# Striatal FOXP2 is essential for stable vocal production in non-human primates

**DOI:** 10.64898/2026.05.06.723125

**Authors:** Hailin Liu, Yuting Yao, Chen Wang, Xiaozhi Sun, Yaolin Zhang, Kaiyi Liu, Rongrong Yang, Lili Zhang, Liangtang Chang, Chunlong Xu, Junfeng Huang, Neng Gong

## Abstract

The transcription factor *FOXP2* is the most well-known language-related gene in humans, yet its role in primate vocalization remains poorly understood. Here we report that knockdown of FOXP2 in the striatum markedly disrupts vocalization stability in the marmoset monkey, a valuable non-human primate model for studying vocal behavior. FOXP2 exhibited high expression in the marmoset striatum, especially during early development. Using the CRISPR-Cas12 system, we achieved specific *in vivo* editing of the *FOXP2* gene and effective knockdown of FOXP2 protein expression in the marmoset striatum. Two neonatal marmosets received bilateral striatal injections of the gene-editing and control virus, respectively, and were raised together in the same family. In three such marmoset pairs, analysis of vocalizations recorded during 6–15 weeks post-injection revealed that striatal FOXP2 knockdown significantly altered vocal features and increased intra-individual variability in phee syllables—the most common marmoset vocalization, often produced repetitively as multi-syllable phee calls. Notably, in *FOXP2*-edited marmosets, acoustic alterations were minimal in the first syllable of phee calls but became progressively more pronounced in subsequent syllables, which exhibited a marked upward shift in the frequency spectrum over time with progressively steeper slopes. These temporal dynamics in vocal features reflect a reduction in the stability of continuous vocal production. In line with the known striatal functions in motor control, our findings provide the first evidence of *FOXP2* in controlling vocalization in non-human primates, thereby opening new avenues for investigating the neural mechanisms underlying *FOXP2* function.

## Introduction

Language, a defining human trait that underpins communication and complex cognition, stands as a distinctive outcome of human evolution and development^1^. Nevertheless, its genetic foundations remain largely elusive. *FOXP2*, the first gene found to be related to language, was identified through studies of the “KE family,” which exhibits an inherited speech and language disorder^2^. Affected individuals carrying *FOXP2* mutations typically display pronounced developmental speech deficits^3^. Evolutionarily, *FOXP2* is not unique to humans but is relatively conserved across species^4^. In diverse vertebrates including mammals, birds, and fish, its gene sequence and expression patterns show remarkable similarities^5–7^. Notably, the human FOXP2 protein differs from that of chimpanzees and macaques by only two amino acids^8,9^. While the functional implications of these subtle differences for the evolution of human language remain unclear^7,10^, such strong evolutionary conservation suggests that *FOXP2* contributes to the emergence of language through ancient molecular pathways and enables the functional investigation of *FOXP2* in non-human animal models^1^.

A substantial body of evidence underscores the critical role of the basal ganglia in language function. As a central structure within the basal ganglia, the striatum exhibits consistently high levels of FOXP2 expression across species^6,11–14^. In humans, disruptions in *FOXP2* lead to language impairments accompanied by functional abnormalities in the striatum^15^. Mouse models further corroborate this association: both *Foxp2*-R552H knock-in mice carrying the mutation identified in the KE family and mice with the humanized *Foxp2* allele display altered striatal activity^16,17^. Additionally, striatal knockdown of Foxp2 in mice results in Huntington’s disease–like motor deficits, whereas overexpression of Foxp2 in the striatum of Huntington’s disease model mice alleviates motor symptoms^18^. Songbirds, a well-established model for vocal learning and production, possess a specialized region in the basal ganglia known as Area X—a core node of the song system that shares cellular composition and connectivity features with the mammalian striatum^19–21^. In Area X, FoxP2 expression fluctuates dynamically with vocal behavior^22,23^, and both knockdown and upregulation of FoxP2 in this region impairs vocal performance and learning in songbirds^19,24^. These findings underscore the pivotal role of striatal *FOXP2* in regulating vocal behavior and motor control.

Despite existing research on *Foxp2/FoxP2* in rodents and birds, the role of the *FOXP2* gene in human speech and vocal development remains poorly understood. These limitations are compounded by the substantial evolutionary divergence between rodents/birds and humans. Notably, functional studies investigating the role of *FOXP2* in vocal behavior have yet to be conducted in primates. The common marmoset, a small New World monkey, has recently gained prominence as a valuable non-human primate model for vocal communication research^25,26^. Similar to songbirds, marmosets exhibit a diverse repertoire of context-dependent call types^27,28^. Furthermore, vocal production in infant marmosets undergoes pronounced developmental changes^29,30^, with the maturation of their vocal behavior dependent on both intrinsic physiological development and social vocal interactions with parents^31–33^. The FOXP2 protein in marmosets shows high sequence conservation with that of Old World monkeys such as macaques, differing by only a single amino acid^8^. Thus, the rich vocal repertoire and dynamic developmental trajectory of vocalizations in marmosets offer a compelling primate model for elucidating the function of *FOXP2* in vocal development and production. In this study, we characterized developmental changes of FOXP2 expression within the marmoset striatum, utilized gene-editing approaches to achieve targeted knockdown of striatal FOXP2 during early development, and evaluated the consequent effects on vocal development and production in marmosets.

## Results

### Expression patterns of FOXP2 in the marmoset striatum

We first examined the expression patterns of FOXP2 in the marmoset brain. Immunohistochemical analysis of brain sections from one infant (7 weeks old) and one adult (2 years old) marmoset revealed that FOXP2 was strongly expressed in both the caudate nucleus (Cd) and putamen (Pu) of the striatum, consistent with previous reports^34,35^ (Fig. 1a–c). In cortical regions, FOXP2 expression was mainly localized to deep layers IV–VI (Fig. 1b). Quantification of FOXP2-positive cells showed that striatal expression was higher in the infant marmoset compared to the adult (Fig. 1c). To corroborate these findings, we performed RNAscope to assess the distribution of *FOXP2* mRNA. RNA expression analysis confirmed a relatively high proportion of *FOXP2*-positive cells in both the striatum and deep cortical layers, with striatal expression remaining significantly higher in the infant than in the adult (Fig. 1d, e). Together, these results indicate that FOXP2 is highly expressed throughout the striatum in marmosets, particularly during early developmental stages.

**Fig. 1.**
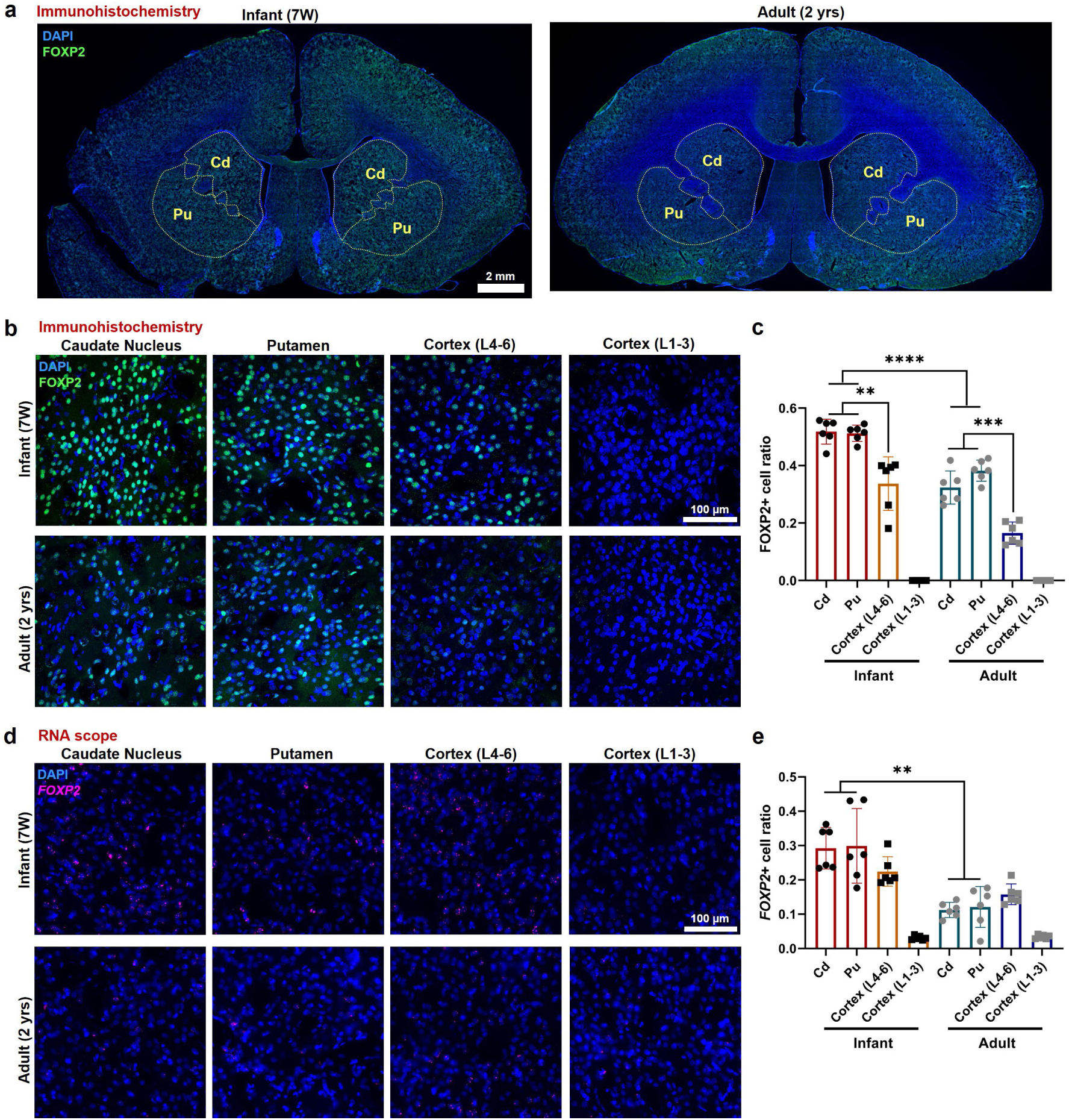
Expression patterns of FOXP2 in infant and adult marmoset brains. **a**, Representative images showing FOXP2 expression in the infant and adult marmoset brains. Cd, Caudate nucleus. Pu, putamen. **b**, Representative images showing FOXP2 expression in the striatum and cortex of infant and adult marmoset brains. **c**, The FOXP2+ cell ratio in the striatum and cortex of infant and adult marmoset brains in immunohistochemical analysis. Data are mean ± s.d.; \*\**P* < 0.01; \*\*\**P* < 0.001; \*\*\*\**P* < 0.0001; unpaired *t*-tests. **d**, Representative images showing *FOXP2* RNA distribution in the striatum and cortex of infant and adult marmoset brains. **e**, The *FOXP2*+ cell ratio in the striatum and cortex of infant and adult marmoset brains in RNAscope analysis. Data are mean ± s.d.; \*\**P* < 0.01; unpaired *t*-tests.

### Gene editing of *FOXP2* in the marmoset striatum

To investigate the role of high FOXP2 expression in the striatum during early development, we sought to specifically reduce FOXP2 levels in the marmoset striatum by *in vivo* editing of the *FOXP2* gene at an early developmental stage. We utilized a compact CRISPR system based on Cas12^36^ and identified two guide RNAs with the highest editing efficiency via *in vitro* screening (Fig. 2b and Extended Data Fig. 1). To assess the *in vivo* efficacy of this approach, we injected the Cas12/gRNA construct packaged in AAV9 viral vectors into the unilateral striatum of an adult marmoset. A GFP-expressing virus was co-injected to mark the injection site (Fig. 2a). At 6 weeks post-injection, we evaluated FOXP2 protein expression by immunohistochemistry and found that the proportion of FOXP2-positive cells in the injected striatum was significantly lower than that on the contralateral uninjected side with a reduction of more than half (Fig. 2c–e). Moreover, sequencing of tissue from the injection site confirmed deletions near the guide RNA target sites within the *FOXP2* gene, validating successful gene editing (Fig. 2f and Extended Data Fig. 2a, c).

**Fig. 2.**
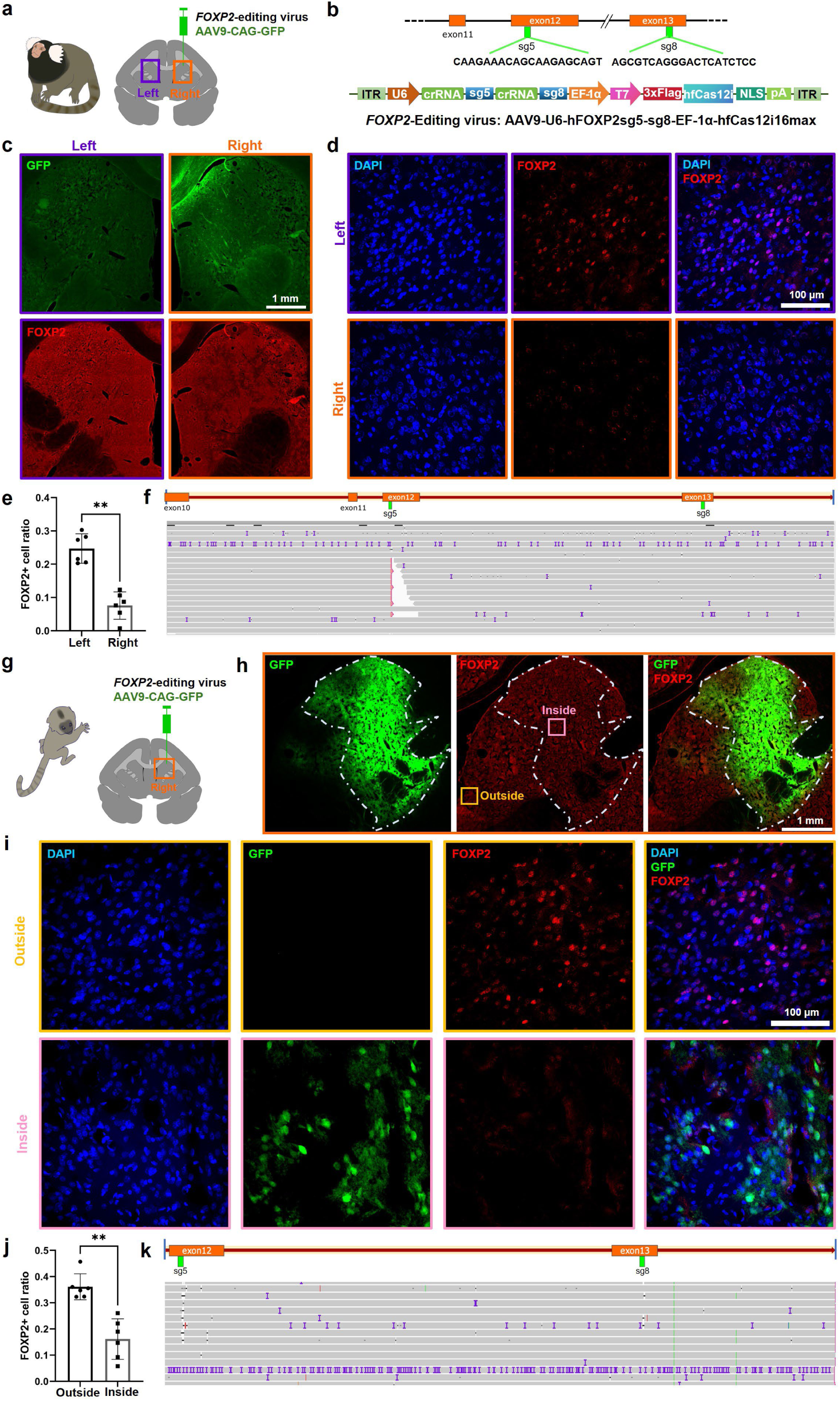
Gene editing of *FOXP2* in the marmoset striatum. **a**, The editing virus and AAV9-GFP were mixed and injected into the right striatum of an adult marmoset. **b**, Schematic diagram of virus vector construction for *FOXP2* editing. The two sgRNAs, sg5 and sg8, were targeting exon12 and exon13 respectively. **c**, Representative images showing GFP and FOXP2 signals in the striatum. Images on the left were from the non-injected side, and images on the right were from the injected side. **d**, Representative images showing FOXP2+ cells in the striatum. Images on the top were from the non-injected side, and images on the bottom were from the injected side. **e**, The FOXP2+ cell ratio in left and right caudate nucleus. Data are mean ± s.d.; \*\**P* < 0.01; unpaired *t*-tests. **f**, The result of third generation sequencing (TGS) of the tissue from the injection side. The purple “I”s refer to mismatches. Blanks refers to deletions. The frequencies for all deletion sizes are presented in Extended Data Fig. 2. **g**, The editing virus and AAV9-GFP were mixed and injected into the right striatum of an infant marmoset. **h**, Representative images showing GFP and FOXP2 signals of right caudate nucleus. GFP signal represents the infected area, which was circled by white dashed line. **i**, Representative images showing FOXP2+ cells inside and outside the infected area. Images on the top were from the injected area, and images on the bottom were outside the injected side. Scale bar, 100 μm. **j**, The FOXP2+ cell ratio of inside and outside the infected area. Data are mean ± s.d.; \*\**P* < 0.01; unpaired *t*-tests. **k**, TGS result of the tissue from the injection side. The purple “I”s refer to mismatches. Blanks refers to deletions. The frequencies for all deletion sizes are presented in Extended Data Fig. 2.

We next evaluated the efficacy of the Cas12 system for editing the *FOXP2* gene in the striatum of infant marmoset by co-injecting the same gene-editing virus along with a GFP-expressing virus into the unilateral striatum of an infant marmoset at the age of 4 weeks (Fig. 2g). Seven weeks post-injection, immunohistochemistry and gene sequencing were performed at the injection site. GFP expression clearly delineated the infected area, enabling direct comparison of FOXP2-positive cells in infected versus uninfected regions within the same brain section (Fig. 2h). We again observed an approximately 50% reduction of FOXP2-positive cells within the infected region relative to the uninfected area (Fig. 2i, j). Sequencing results further confirmed deletions in the *FOXP2* gene near the guide RNA target sites (Fig. 2k and Extended Data Figs. 2b, c). Together, these results demonstrate that the Cas12 system effectively edits the *FOXP2* gene and reduces its expression in the striatum of both infant and adult marmosets by about half, a level sufficient to impair FOXP2 function in the striatum.

### Striatal FOXP2 knockdown in neonatal marmosets alters vocal features

Based on this effective editing strategy, we used the Cas12 system to target the *FOXP2* gene specifically in the striatum of neonatal marmosets and assessed the consequent behavioral phenotypes. Four pairs of neonatal marmosets were selected, including three twin pairs and one non-twin pair (Extended Data Table 1). At approximately two weeks after birth, one marmoset from each pair received bilateral injections of a *FOXP2*-editing virus into the striatum, while its counterpart was injected with a control virus in the same region, with three injection sites per hemisphere per animal (Fig. 3a). After the procedure, each pair was hand-reared for 1–3 days until cranial wounds had largely healed, after which they were returned to a family with normal parental care (Fig. 3b). From 6 to 15 weeks post-injection, vocalizations were recorded weekly from both *FOXP2*-edited and control marmosets. Each animal was individually placed in a sound-proof chamber and recorded for 30 minutes (Fig. 3b). Among the four pairs, three produced a sufficient number of calls for quantitative analysis. In the remaining pair, neither the *FOXP2*-edited nor the control animal vocalized during recordings. We thus subsequently collected striatal tissue samples from this pair and performed sequencing analysis, which confirmed substantial editing of the *FOXP2* gene in the striatum of the gene-edited marmoset, whereas the *FOXP2* sequence in the control animal remained comparable to that of wild-type marmosets (Extended Data Fig. 3). These results further validated the striatal *FOXP2* editing and subsequent FOXP2 knockdown in neonatal marmosets. Additionally, the *FOXP2*-edited marmosets exhibited generally normal physical development, with no significant differences in body weight between the *FOXP2*-edited and control groups (Extended Data Fig. 4). Behavioral observations during vocal recordings also revealed no detectable abnormalities of global motor ability in the *FOXP2*-edited animals (Extended Data Fig. 5).

**Figure 3.**
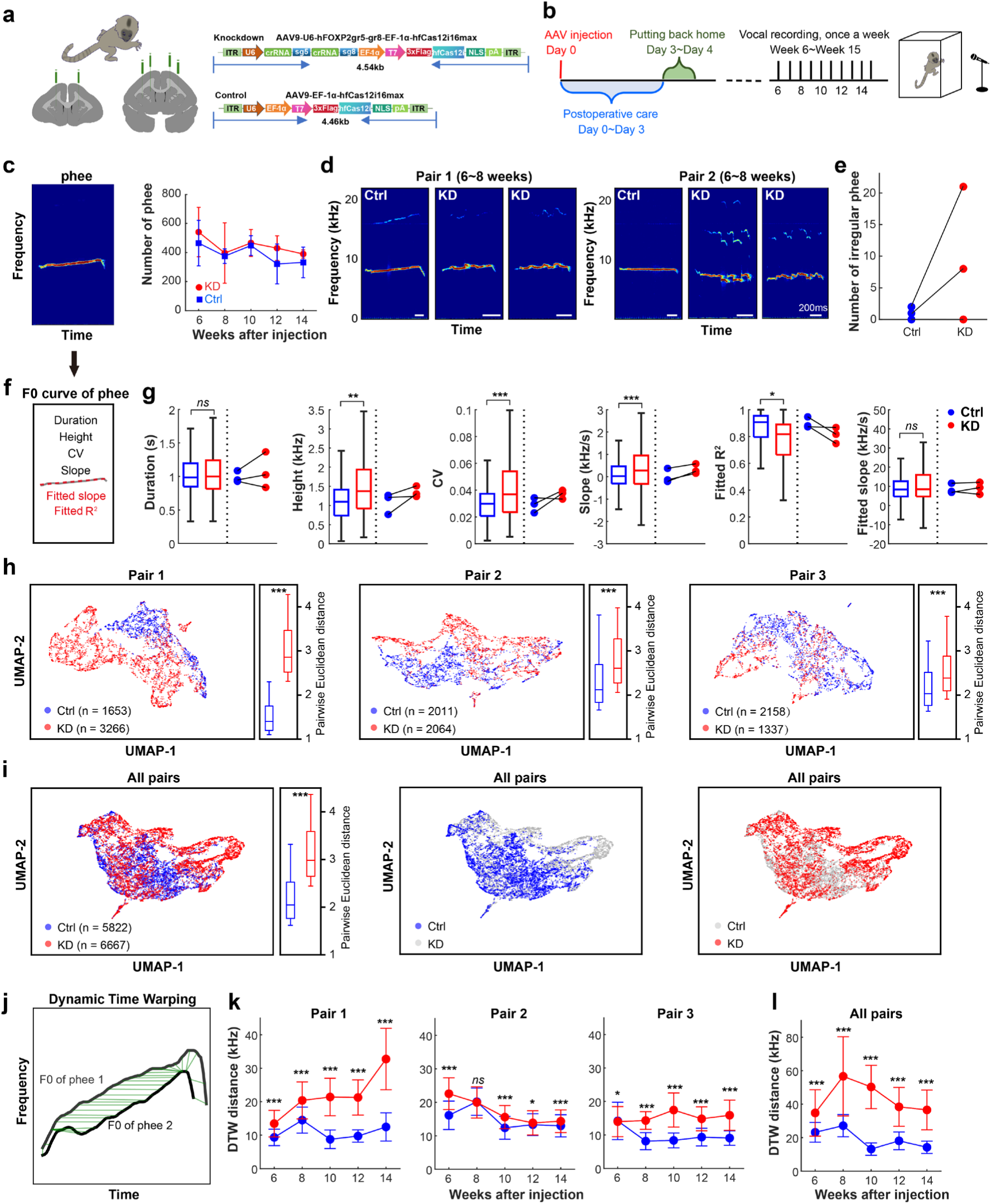
Striatal FOXP2 knockdown alters vocal features and reduces intra-individual vocal similarity. a, Each neonatal marmoset received bilateral injections of a *FOXP2*-editing virus or a control virus into the striatum, with three injection sites per hemisphere. **b,** Each animal was hand-reared for 1–3 days after virus injection, and then returned to a family with normal parental care. Vocal recordings were performed once per week from 6 to 15 weeks post-injection. **c,** Representative spectrogram of a typical phee syllable, and the total number of phee syllables in the control (Ctrl) and knockdown (KD) groups during the 6–15 weeks post-injection period. **d,** Representative spectrograms of irregular phee syllables. **e,** Number of irregular phee syllables in the Ctrl and KD groups. Lines indicate marmosets from the same pair. **f,** Schematic illustration showing the fundamental frequency (F0) curve of a phee syllable and its shape characteristics. **g,** Comparison of F0 features between the Ctrl and KD groups. For each subplot, the left panel shows box plots indicating the 25th, 50th, and 75th percentiles, with whiskers extending to 1.5 × the interquartile range (IQR); statistical significance was assessed using linear mixed-effects models. The right panel shows the median value for each individual marmoset from the three pairs. **h,** For each marmoset pair, the left panel displays the distribution of phee syllables in a two-dimensional UMAP space by the six F0 curve features. The right panel shows intra-individual Euclidean distances between syllables in the original six-dimensional feature space; *P* values were calculated using two-sided permutation tests. **i,** The UMAP distribution of phee syllables from all Ctrl and KD animals and the corresponding intra-individual Euclidean distances between syllables. **j,** Schematic illustration showing the calculation of dissimilarity between two phee syllables using dynamic time warping (DTW) applied to F0 curves. **k** and **l,** Intra-individual phee syllable DTW distances for each pair and all pairs. Data are median ± MAD. *P* values were calculated using the Mann–Whitney U test. *ns*, not significant (*P* > 0.05), \**P* < 0.05, \*\**P* < 0.01, \*\*\**P* < 0.001.

Under the isolated condition, the majority of vocalizations captured were “phee” calls (Fig. 3c), a typical long-distance contact call in marmosets^28,37^. In the three successfully recorded marmoset pairs, phee calls collected from two consecutive weeks were combined into a single time point for analysis. No significant differences were found in the number of phee syllables produced between the *FOXP2*-edited and control groups during the 6–15 weeks post-injection developmental period (Fig. 3c). However, at the first recording time point (6–8 weeks), two of the three *FOXP2*-edited marmosets produced phee calls with abnormal waveforms, characterized by vocal discontinuities and tremors (Fig. 3d, e). Since this period is critical for vocal development in marmosets, the downregulation of FOXP2 protein in the striatum may have disrupted the normal process of vocal maturation.

The peak frequency of phee syllables produced by *FOXP2*-edited marmosets showed no significant difference compared to the control group (Extended Data Fig. 6). We subsequently extracted the fundamental frequency (F0) curve of each phee syllable and analyzed its shape characteristics, including duration, height, coefficient of variation (CV), slope, as well as linearity and slope of a linear fit to the F0 curve (Fig. 3f). Compared with controls, the *FOXP2*-edited marmosets exhibited significantly greater height, slope, and CV, along with reduced fit linearity in the F0 curve (Fig. 3g). These differences were consistently observed across all three marmoset pairs (Fig. 3g). The increased CV and reduced fit linearity of F0 curves in *FOXP2*-edited animals suggest reduced vocal stability in their phee call production. Thus, FOXP2 knockdown in the striatum of neonatal marmosets shows specific effects on vocal maturation and production, with remarkable alterations in the vocal features.

### Striatal FOXP2 knockdown reduces intra-individual vocal similarities

We then further examined the stability of vocalization by quantifying intra-individual similarity among phee syllables. After projecting phee syllables into a UMAP space based on six F0 characteristics, UMAP visualization revealed that *FOXP2*-edited marmosets exhibited a more dispersed distribution of phee syllables than controls (Fig. 3h, i). This pattern was consistent both within individual pairs and across the full dataset (Fig. 3h, i). Analysis of the Euclidean distance, calculated from raw F0 characteristics between syllables, also showed that *FOXP2*-edited marmosets exhibited significantly larger intra-individual distances than controls, in both pairwise and overall comparisons (Fig. 3h, i). We also employed dynamic time warping (DTW) to directly compare the similarity of F0 curves among all syllables produced by each marmoset (Fig. 3j).

Longitudinal analysis revealed a significant reduction in the intra-individual similarity of F0 curves in *FOXP2*-edited marmosets compared to controls during 6–15 weeks after injection. This effect was consistently observed within each animal pair (Fig. 3k) and was also significant across the full dataset (Fig. 3l). Moreover, this difference in intra-individual vocal similarity between *FOXP2*-edited and control animals tended to increase over time (Fig. 3k, l). Together, these findings demonstrate that *FOXP2*-edited marmosets exhibit significantly lower intra-individual similarity in phee call production, indicating diminished vocal stability following striatal FOXP2 knockdown.

### Striatal FOXP2 knockdown reduces stability in continuous vocal production

Marmoset phee syllables often occur sequentially, forming multi-syllable phee calls^38^. Examination of the spectrograms across all three marmoset pairs revealed a consistent pattern: while the first phee syllable in multi-syllable calls showed little difference between groups, subsequent syllables produced by *FOXP2*-edited marmosets exhibited progressively steeper F0 trajectories (Fig. 4a). This visually discernible divergence may contribute to the reduced vocal similarity observed in the *FOXP2*-edited group (Fig. 3). To further investigate this pattern, we color-coded the UMAP projection of all phee syllables from Fig. 3i based on their sequential position within calls (Fig. 4b). This UMAP visualization showed substantial overlap between groups for syllables in the 1^st^ position, whereas syllables in the 2^nd^ and 3^rd^ positions displayed clear separation in UMAP space (Fig. 4b). To quantify this observation, we calculated the UMAP centroid for syllable clusters at each sequence position. The resultant inter-centroid distances between the *FOXP2*-edited and control groups demonstrated that vocal divergence increases progressively throughout the syllable sequence (Fig. 4c).

**Fig. 4.**
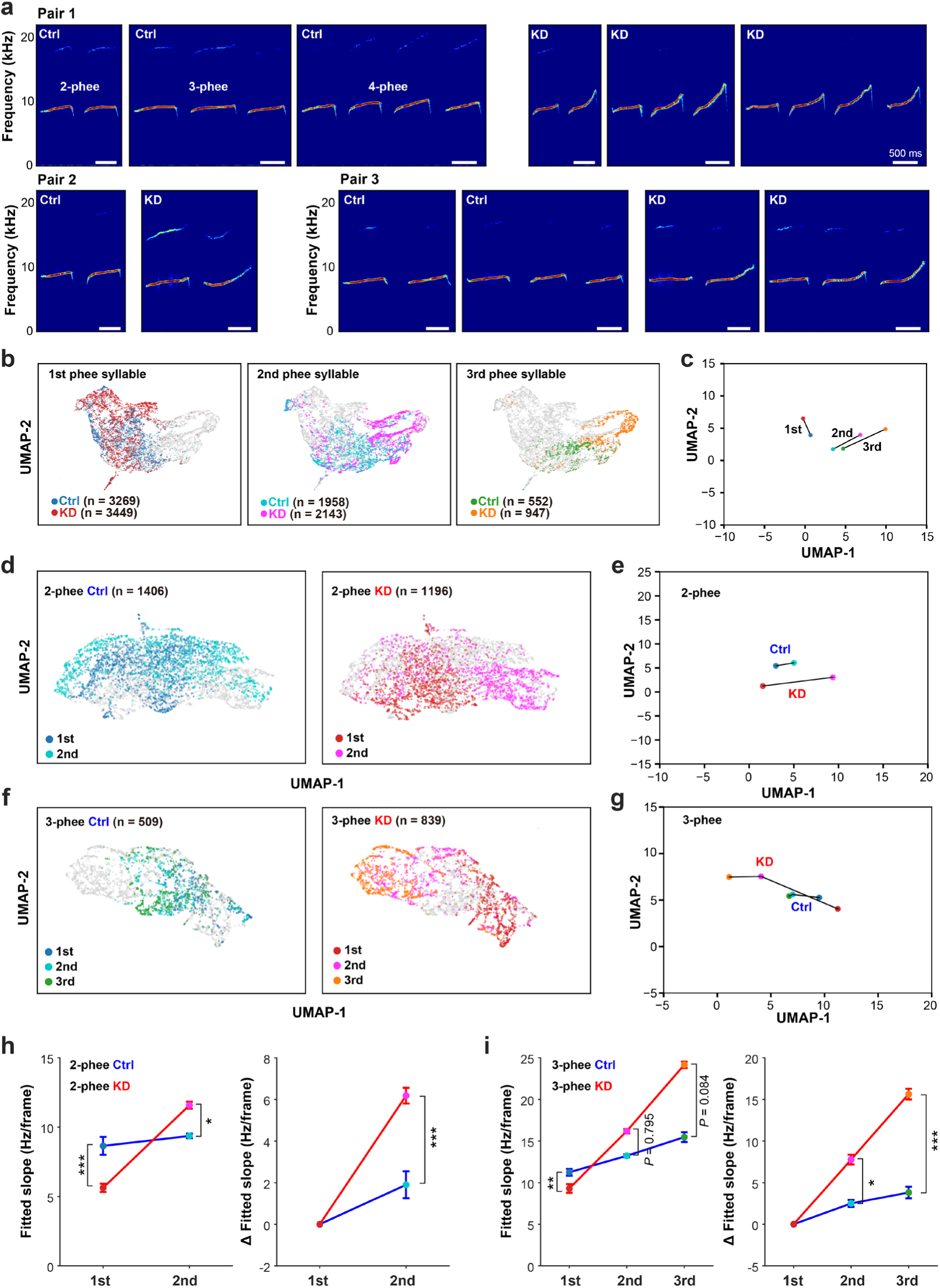
Striatal FOXP2 knockdown reduces stability in continuous vocal production. a, Representative spectrograms of multi-syllable phee calls from three pairs. **b,** Recoloring phee syllables at different positions for the control (Ctrl) and knockdown (KD) groups in the same UMAP space shown in Fig. 3i; *n* denotes the number of syllables. **c,** Median positions of the 1st, 2nd, and 3rd phee syllables for the Ctrl and KD groups shown in **b**. **d,** UMAP distribution of the 1st and 2nd phee syllables from 2-phee calls in the Ctrl (left panel) and KD (right panel) groups. **e,** Median positions of the 1st and 2nd phee syllables shown in **d**. **f** and **g,** UMAP distribution of the 1st, 2nd, and 3rd phee syllables from 3-phee calls in the Ctrl and KD groups. **h,** Left, fitted slopes of the 1st and 2nd phee syllables from 2-phee calls in the Ctrl and KD groups. Right, fitted slopes of the 2nd phee syllable normalized to the 1st syllable, calculated by subtracting the fitted slope of the 1st syllable. **i,** Left, fitted slopes of the 1st, 2nd, and 3rd phee syllables from 3-phee calls in the Ctrl and KD groups. Right, fitted slopes of the 2nd and 3rd syllables after normalization to the 1st syllable. Statistical significance was assessed using linear mixed-effects models. \**P* < 0.05, \*\**P* < 0.01, \*\*\**P* < 0.001.

We then focused our analysis on the most common types of multi-syllable phee calls—specifically, double-syllable (2-phee) and triple-syllable (3-phee) calls. For all such calls produced by both *FOXP2*-edited and control marmosets, we applied UMAP-based dimensionality reduction to 2-phee and 3-phee calls, respectively. For both phee call types, syllables from different sequence positions exhibited clearer separation in UMAP space in the *FOXP2*-edited group compared to controls; this was demonstrated by significantly greater inter-centroid distances (Fig. 4d–g). To quantify such progressively steeper waveform profiles, we examined the fitted slope of the F0 curve for each phee syllable in 2-phee and 3-phee calls, respectively. We found that in *FOXP2*-edited animals, the F0 slope increased substantially across sequential syllable positions, a trend that was markedly more pronounced than in controls (Fig. 4h, i). Together, *FOXP2*-edited marmosets exhibit a progressive decline in vocal stability during continuous vocalizations, whereas control animals maintain relatively consistent vocal production across sequential syllables, indicating that striatal FOXP2 knockdown reduces stability in continuous vocal production in marmosets.

## Discussion

Since its discovery, the *FOXP2* gene has been extensively investigated in animal models such as mice and songbirds, including studies involving Foxp2/FoxP2 knockout, overexpression, and knock-in of the mutated or humanized form^1,17,19,24^. However, no studies have yet demonstrated the role of *FOXP2* in regulating vocal behavior in primates. The commonly used macaque model exhibits limited vocal capabilities and is thought to lack vocal learning ability^25,39^. The emergence of the marmoset as a model offers a promising alternative, as marmosets display diverse call types, engage in vocal communication, and undergo vocal development processes^29,30,32^. Another major limitation has been the difficulty of editing the *FOXP2* gene in developing primates. *FOXP2* is critical for motor functions not limited to vocal production. Embryonic knockout of Foxp2 in mice could cause severe motor impairments^40^ and result in early lethality^41^. Thus, embryonic knockout of FOXP2 in primate models is not feasible. In this study, we established a method for localized gene editing within specific brain regions of neonatal marmosets. Surgical damage was minimized, and following virus injection, infant marmosets were hand-reared for several days until wound healing. Vocal development in marmosets is highly dependent on the family environment and parental feedback^31–33^, and individuals from the same family typically exhibit similar call features^42^, thus we used twin marmosets or paired marmosets raised in the same family for this study, with one receiving the editing virus while the other was injected with a control virus. Such paired marmosets were returned to the same family for rearing under normal parental care, enabling reliable analysis of vocal phenotypes by comparing the gene-edited marmoset and its littermate control. For viral delivery, we have screened multiple AAV subtypes for infection efficiency in the marmoset brain and identified AAV9 as the most effective for localized brain injections^43^. In contrast to the conventional Cas9 system, we employed the more compact Cas12 system^44^ for gene editing in marmosets. This allowed both the editing machinery and gRNA to be packaged into a single virus, thereby improving delivery and editing efficiency in the marmoset brain. Together, we have developed an experimental approach that enables localized gene editing in the brains of neonatal marmosets, thus providing a valuable approach for studying gene function during early development in marmoset models.

Previous studies in humans, mice, and songbirds have consistently implicated the basal ganglia as a key region through which *FOXP2* influences vocal behavior^1,4,45^. Consistent with these findings, we observed high expression of FOXP2 in the marmoset striatum, a central component of the basal ganglia, particularly during early developmental stages. We therefore targeted *FOXP2* in the striatum of neonatal marmosets for gene editing. Following successful reduction of FOXP2 protein levels in the striatum of neonatal marmosets, the edited animals exhibited nearly normal physical development and general motor activity. However, they showed specific alterations in vocal behavior. In particular, marmosets with striatal FOXP2 knockdown displayed reduced stability in continuous vocal production, a finding consistent with the known role of the striatum in fine motor and sequential movement control^46,47^. Notably, the vocal instability observed in these *FOXP2*-edited animals resembles certain clinical features reported in patients carrying *FOXP2* mutations, who present with speech disorders characterized by shortened utterances, articulation errors, and impaired sequential orofacial movements^48^. Our study demonstrates that endogenous FOXP2 in the marmoset striatum is essential for the production of stable vocalizations. Future studies by introducing and expressing a humanized *FOXP2*—carrying the two human-specific amino acid changes—in marmosets could substantially advance our understanding of the functional contributions of these residues to vocal behavior and provide important insights into the evolution of human language.

In the present study, we did not explore the neural mechanisms through which striatal FOXP2 knockdown leads to impaired vocal stability in marmosets. However, given the evolutionary conservation of *FOXP2* and basal ganglia circuitry across species^4^, relevant insights can be drawn from studies in mice and songbirds. *Foxp2/FoxP2* is also highly expressed in the mouse striatum^12^ and in Area X, a basal ganglia region in songbirds^11^. In mice, introduction of the *Foxp2*-R552H mutation identified in the KE family disrupts striatal circuit function during motor skill learning, manifesting as reduced neuronal firing frequency and impaired temporal coordination of striatal activity^16^. Mice carrying a humanized form of *Foxp2* show altered ultrasonic vocalizations and reduced exploratory behavior^17,49^, accompanied by decreased dopamine levels in the brain and increased dendritic length of striatal neurons^17^. Furthermore, knockdown of Foxp2 in the striatum of wild-type mice induces Huntington’s disease-like symptoms, whereas overexpression of Foxp2 in a Huntington’s disease mouse model alleviates motor deficits and alters the expression of synaptic function-related genes^18^. Songbirds, unlike mice, are more widely used as animal models for vocal behavior and language-related research^50^. Area X serves as a central node in the songbird vocal pathway and belongs to the avian basal ganglia^19^.

Perturbations of FoxP2 levels in Area X by either knockdown or overexpression impair vocal imitation and learning in songbirds^19,24^. Moreover, FoxP2 expression in Area X is dynamically regulated by vocal behavior: it decreases when males sing alone but remains stable when they sing to females^22^. In hearing-intact zebra finches, FoxP2 levels progressively decline with prolonged singing, an effect absent in deafened birds^23^. Mechanistic studies suggest that reduced FoxP2 expression in Area X disrupts the balance of dopamine receptor expression between the striatal direct and indirect pathways, leading to aberrant syllable sequencing^11^. Based on these findings, we speculate that FOXP2 knockdown in the marmoset striatum may similarly impair cortico-basal ganglia circuits and dopaminergic pathways, thereby compromising stable vocal production.

In this study, we successfully achieved specific editing of the *FOXP2* gene and significant downregulation of FOXP2 protein expression within the striatum of neonatal marmosets, and demonstrated that striatal FOXP2 is essential for stable vocalization, especially continuous vocal production in marmosets. Such striatal FOXP2 knockdown in marmosets effectively recapitulated key speech disorders seen in patients carrying *FOXP2* mutations, underscoring the value of the marmoset as a non-human primate model for probing the mechanisms of human language and the function of language-related genes. Furthermore, our results support the hypothesis that *FOXP2*, a pivotal gene for human language, contributed to the evolution of human language through its deeply conserved role in regulating vocal behavior in primates.

## Acknowledgements

We thank Mu-ming Poo for discussions and comments on this manuscript, and the marmoset facility at CAS Center for Excellence in Brain Science and Intelligence Technology for assistance in animal surgery and sound recording. **Funding:** This work was supported by Brain Science and Brain-like Intelligence Technology – National Science and Technology Major Project 2021ZD0203900; Shanghai Municipal Science and Technology Major Project “The pathogenesis of Alzheimer’s disease and the development of related innovative drugs”, Project 22ZR1481500 and Eastern Talent Plan Project; CAS “Strategic Priority Research Program” (XDB1010101); NSFC Project 32371085; National High-level Young Top Talent Support Program (E519202A02), Shanghai Rising Star Program for Science and Technology (22QA1412300), Shanghai Oriental Talent Program (E544201A02) and intramural funds of Lin-Gang Laboratory (LG-GG-202402-01, LGL-0121, LG-GG-202401-01-ADA060101, LG-GG-202401-01-ADA070102, LGL-3142-ADB130100, PDA010100, 1100000A01).

## Author contributions

N.G. conceived and designed the study. H.L., Y.Y., C.W., X.S., K.L., R.Y., L.Z. and C.X. performed animal surgery, gene editing, molecular and biochemical analyses. C.W., Y.Z., L.C. and J.H. performed sound recordings and analyses. Y.Y., X.S., C.X., J.H. and N.G. prepared the figures. N.G., Y.Y. and C.X. wrote the manuscript. All authors have read and approved the manuscript submission.

## Data availability

The data that support the findings of this study are available from the corresponding author upon request.

## Code availability

MATLAB scripts in this study are available from the corresponding author upon request.

## Methods

### Subjects

Twelve marmosets (*Callithrix jacchus*), including 10 infants and 2 adults, were used in this study. Animals were maintained in a temperature and humidity-controlled facility on a 12-h (07:00–19:00) light/dark (LD) cycle, with ad libitum access to food and water until experimental use. All procedures were approved by the Animal Care and Use Committee of the Institute of Neuroscience, Center for Excellence in Brain Science and Intelligence Technology, Chinese Academy of Sciences.

### Immunohistochemistry staining and RNAscope

The marmosets were perfused with 4% paraformaldehyde (PFA), and the brains were taken out and fixed in 4% PFA for 12 h. After 4 days of dehydration in 30% sucrose/ phosphate-buffered saline (PBS), the brains were embedded in Optimal Cutting Temperature (OCT) compound at -20°C and cut on a freezing microtome (Cryostar NX50, Thermo Scientific, MA, USA) at 30 µm per section. The sections were immersed in PBS to wash off the OCT compound, then flattened on slides and heated to 54°C for 54 min to be fixed on the slides. After 10 minutes of antigen retrieval with 0.05% trypsin at RT, the sections were blocked in blocking buffer (PBS containing 0.3% Triton X-100 and 5% bovine serum albumin) for 2 h at RT. The sections were incubated with rabbit anti-FOXP2 (1:500, Cambridge, UK) at 4°C overnight and secondary antibodies (donkey-anti-rabbit-488 or donkey-anti-rabbit-594, 1:600, Invitrogen, CA, USA ) at RT for 4 h. DAPI (1:400, Solarbio, Beijing, China, MO) was used to stain nuclei. Images were acquired on an Olympus VS120 Slide Scanning System (Olympus, Tokyo, Japan) and a Nikon NiE-A1 plus confocal microscope (Nikon, Tokyo, Japan) and analyzed by ImageJ (NIH, Bethesda, USA). For the data analysis, the signals of each channel were first identified by Cellpose and were corrected manually.

The experimental procedures of RNAscope were mostly followed the official instructions (https://acdbio.com/documents/product-documents). The brain sections were incubated with 3% H_2_O_2_-PBS solution for nearly 1 hour until the sections no longer produced bubbles after the OCT-washing-off in PBS. Then the sections were fixed in 4% PFA at 4°C for 15 minutes, followed by 5 minutes of target retrieval at 99°C in RNAscope® Target Retrieval Reagents. After washing off in PBS, the sections were flattened on slides and heated to 54°C for 54 min to be fixed on the slides. The next steps begin with applying RNAscope® Protease 3 and then follow the instructions. The fluorophore used was TSA® Plus Cyanine 5 (1:1000, Akoya Biosciences, MA, USA).

Images were acquired on an Olympus VS120 Slide Scanning System (Olympus, Tokyo, Japan) and a Nikon NiE-A1 plus confocal microscope (Nikon, Tokyo, Japan) and analyzed by ImageJ (NIH, Bethesda, USA). For the data analysis, the signals were first identified by Cellpose and were corrected manually.

### Gene editing

To identify efficient single-guide RNAs (sgRNAs), candidates were designed adhering to the 5’-TTN-3’ Protospacer Adjacent Motif (PAM) requirement of hfCas12iMAX, with a protospacer length of 20 nucleotides (nt). Subsequently, these sgRNA sequences were cloned into a plasmid vector co-expressing the hfCas12iMAX protein. To assess cleavage and editing efficiency in mammalian cells, HEK293T cells were seeded into 24-well plates (Corning) and cultured until reaching approximately 80% confluency. Cells were then transfected with 2 μg of plasmid encoding hfCas12iMAX and the target sgRNA using Polyethyleneimine (PEI, Polysciences). The transfection was performed at a PEI-to-DNA ratio of 2.5:1 (v/w). Forty-eight hours post-transfection, cells were dissociated using 0.25% Trypsin (Gibco). To enrich transfected populations, the top 20% of GFP-positive cells were collected via Fluorescence-Activated Cell Sorting (FACS). The sorted cells were subsequently lysed for genomic DNA extraction and editing analysis. Two effective guide RNAs targeting FOXP2 were identified: sg5 (CAAGAAACAGCAAGAGCAGT) and sg8 (AGCGTCAGGGACTCATCTCC). Recombinant AAV9 vectors were produced in-house, including the gene-editing virus (AAV9-U6-hFOXP2-sg5-sg8-EF1α-hfCas12i16max) and the non-targeting control virus (AAV9-U6-EF1α-hfCas12i16max).

### Surgery and virus injection

The adult marmoset was not fed overnight before surgery and was anesthetized with 3% inhaled isoflurane while maintained with 1-2% inhaled isoflurane. It was placed in a standard stereotaxic apparatus (RWD, Shenzhen, China) after being anesthetized. A heating pad was used to maintain body temperature and monitors were used to monitor vital signs. It received standard postoperative care and was returned to their cage once they showed signs of recovery from anesthesia.

The infant marmosets were not fed for 3 hours but received 0.3 mL of intravenous 5% glucose injection before surgery and were anesthetized and maintained with 1-2% inhaled isoflurane. Then they were placed in a customized stereotaxic apparatus to minimize surgery injury. A heating pad was used to maintain body temperature and monitors were used to monitor vital signs. After surgery, they received 3 days of postoperative care and were carefully returned to families to receive normal parental care.

To verify gene-editing efficiency, the editing virus and the AAV9-GFP virus were mixed at the volume ratio 9:1, and delivered into the striatum through a 10-μL syringe driven by a micro syringe pump at a flow rate of 2 nL/s (adult marmoset: A/P: 4.3 mm; M/L: -3mm; D/V: −7.6mm; infant marmoset: A/P: 6 mm; M/L: -2.7 mm; D/V: −6.4 mm; 900 nL virus per site). For the injections in 4 pairs of neonatal marmosets, the editing or control viruses were injected into bilateral striatum at a flow rate of 2 nL/s.

The coordinates (A/P, M/L, D/V) (unit: millimeters) of the injection sites were (7, 3, - 7), (5, 3.5, -6), (5, 3.5, -8), (7, -3, -7), (5, -3.5, -6) and (5, -3.5, -8) with 800 nL virus per site. Before removal, the needle was allowed to remain in the brain for 10 min after the cessation of the injection. The titer of the viruses was 1.61×10^13^ vg/ml.

### Targeted deep sequencing and indel analysis

To evaluate gene-editing efficiency, genomic DNA (gDNA) was extracted from transfected HEK293T cells or AAV-transduced Callithrix jacchus brain tissues using the TIANamp Genomic DNA Kit (TIANGEN, DP304) following the manufacturer’s protocol. Target loci were amplified using Phanta Max Super-Fidelity DNA Polymerase (Vazyme, P505) with specific primers designed based on the human (GRCh38) or marmoset (calJac3) reference genomes. For HEK293T samples, a two-step PCR strategy was employed to generate libraries for next-generation sequencing (NGS). In the second round of PCR, specific barcodes were incorporated at the 5′ ends of the primers for multiplexing. The resulting amplicons were purified using the Gel Extraction Kit (Omega, D2500) and sequenced on the Illumina HiSeq platform. Raw reads were demultiplexed using Cutadapt (v2.8) based on the unique barcodes. For marmoset brain tissues, long-read sequencing was performed to capture potential large deletions or complex rearrangements. SMRTbell libraries were constructed from the purified PCR amplicons and sequenced on the PacBio Sequel platform. Raw circular consensus sequencing (CCS) reads were converted to FASTQ format, retaining original Phred quality scores. The CCS reads were quality-filtered using fastplong (v0.4.1) with the following criteria: length between 1,000 bp and 4,000 bp, a qualified base Phred threshold of 20, and a minimum mean quality score of 10. The precise editing events (indels) were quantified using CRISPResso2 (v2.3.3) with default parameters. Untreated cells or brain tissues injected with control AAVs (lacking sgRNAs) were processed in parallel to serve as negative controls for background noise subtraction.

### Vocal recording and acoustic feature analysis

The subject was weighed and then transported from their home cages to a recording cage (900 mm × 800 mm × 785 mm) located in a sound-proof room. Vocal recordings were obtained using a microphone (Audio-Technica AT2031, Tokyo, Japan) positioned approximately 500 mm in front of the recording cage and connected to a sound card (OCTA-CAPTURE UA-1010, Shizuoka, Japan), with signals digitized at a sampling rate of 48 kHz. Recordings were conducted for 20 min once a week from weeks 6 to 15 following virus injection. The recording order of marmosets was pseudo-randomized and balanced within each pair. To assess global motor ability in infant marmosets, locomotor activity was also recorded, and movement trajectories, total distance traveled, and movement duration were quantified.

After recording, the onset and offset times of each call were identified, and call types (e.g., phee, twitter, trill) were manually annotated based on visual inspection of spectrograms in Praat software (version 6.4.01) by one experienced technician and independently verified by a second technician. Because the majority of recorded vocalizations were phee calls, which were the only call type present across all recording sessions, subsequent analyses focused on phee calls. Audio signals were filtered using a zero-phase Butterworth band-pass filter (2–20 kHz) to reduce background noise. Peak frequency was defined as the frequency with maximum power in the power spectrum. To characterize the time–frequency structure of phee calls, filtered signals were segmented into frames using a 30-ms Hanning window with a 10-ms frame shift. Then, 6 features describing the shape of the fundamental frequency (F0) contour were extracted: (1) <u>Call duration</u>, the temporal length of the call; (2) <u>F0 height</u>, the difference between the maximum and minimum F0 values within a call; (3) <u>F0 CV</u>, the coefficient of variation of the F0 contour across the entire call; (4) <u>F0 slope</u>, calculated as the difference between F0 values at call onset and offset divided by call duration; (5–6) <u>F0 fitted slope</u> and <u>coefficient of determination (R²)</u>, obtained by fitting a linear regression to the rising portion of the F0 contour from call onset to the point of maximum F0. In case that the maximum F0 value occurred at the first time point, the linear regression was fitted to the entire F0 contour. Linear regression was performed using the *linregress* function from the *scipy.stats* package (Python 3.12.3; SciPy 1.13.1). To quantify progressive changes in F0 curve steepness across syllables in 2-phee and 3-phee calls, fitted slopes of subsequent syllables were expressed relative to the first by subtraction (Δfitted slope). This normalization controlled for inter-call variability in pitch dynamics, enabling direct comparison of relative changes across sequential syllable positions.

### Intra-individual vocal similarity analysis

To assess intra-individual vocal stability, we quantified the similarity among phee calls produced by the same marmoset. All 6 call-shape features were independently standardized using Z-score normalization to ensure comparability across features with different scales, and were further mean-centered by subtracting the corresponding individual-specific mean. After normalization, each phee call was represented as a six-dimensional feature vector 𝐳_𝑖_ = (𝑧_𝑖1_, 𝑧_𝑖2_, … , 𝑧_𝑖6_). For each phee call produced by a given marmoset, the Euclidean distance between that call and every other phee call from the same individual was computed as 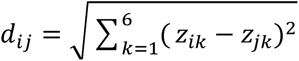 , where 𝑑 *_𝑖j_* denotes the distance between the 𝑖-th and 𝑗𝑗-th phee calls within the same marmoset. The median of these distances, 𝐷_𝑖_ = median *_j_*_≠𝑖_(𝑑*_ij_*), was taken as a measure of call-level dissimilarity. Intra-individual vocal stability was then defined as the mean of 𝐷_𝑖_ across all 𝑁 phee calls produced by each marmoset, 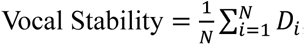 , with lower values indicating higher vocal stability.

To assess whether the difference in intra-individual vocal stability between the two marmosets within each pair was statistically significant, a permutation test with 1,000 iterations was performed. In each iteration, all phee calls from the two individuals were pooled and randomly reassigned into two groups (X and Y), while preserving the original number of calls produced by each marmoset. Vocal stability was computed separately for groups X and Y, and their difference was recorded. The distribution of vocal stability differences obtained from the 1,000 iterations represented the null distribution expected by chance. The observed difference in vocal stability between the two marmosets was compared against this null distribution. If the observed difference exceeded all values obtained from the permutations, the result was considered statistically significant with *P* < 0.001 (Fig. 3h). For intra-group stability in the control (Ctrl) and knockdown (KD) groups, the same vocal stability metric was applied, with phee calls pooled at the group level and individual marmoset identity ignored (Fig. 3i).

To visualize the distribution of F0 call-shape features of phee calls across the Ctrl and KD groups, all six features were first standardized using Z-score normalization. Uniform Manifold Approximation and Projection (UMAP) was then applied to the normalized features of all phee calls to reduce the data to two dimensions. UMAP was only used for visualization and was implemented in Python (version 3.12.3). Parameter values were adjusted slightly to optimize visualization of local versus global structure across different figures. Specifically, for figure 4d, parameters were set to *n_neighbors* = 5 and *min_dist* = 0.5, whereas for the remaining figures, *n_neighbors* = 15 and *min_dist* = 0.1 were used; all other parameters were kept at their default values.

To quantitatively assess dissimilarity between the F0 curves of phee calls, dynamic time warping (DTW) was used to measure distances between time-varying F0 curves. DTW distances were computed using the dtw function in MATLAB R2023a. For each phee call produced by a given marmoset, DTW distances were calculated between that call and all other phee calls from the same individual recorded at the same time point. The median of these pairwise distances was then used as a measure of call-level dissimilarity (Fig. 3k), with larger values indicating greater dissimilarity from other calls. For group-level analyses within the Ctrl or KD group, call-level dissimilarity was computed using the same procedure, with phee calls pooled within each group (Fig. 3l).

### Statistical analysis

All statistical analyses were performed using MATLAB R2023a (MathWorks). Because phee calls from the same individual were not fully independent, acoustic features were analyzed using linear mixed-effects models. Group (Ctrl or KD) was included as a fixed effect, with random intercepts specified for Pair (Pairs 1–3) and for individual marmosets nested within Pair (1 ∣ 𝑃𝑎𝑖𝑟) + (1 ∣ 𝑃𝑎𝑖𝑟: 𝑀𝑎𝑟𝑚𝑜𝑠𝑒𝑡) . For parametric comparisons, the unpaired t-test was used for independent samples, and the paired t-test was used for paired samples. For nonparametric comparisons, the Mann–Whitney U test was used for independent samples, and the Wilcoxon signed-rank test was used for paired samples. Permutation tests were used to assess differences in vocal similarity between Ctrl and KD groups. All tests were two-tailed unless otherwise specified.

## Extended Data

**Extended Data Fig. 1.**
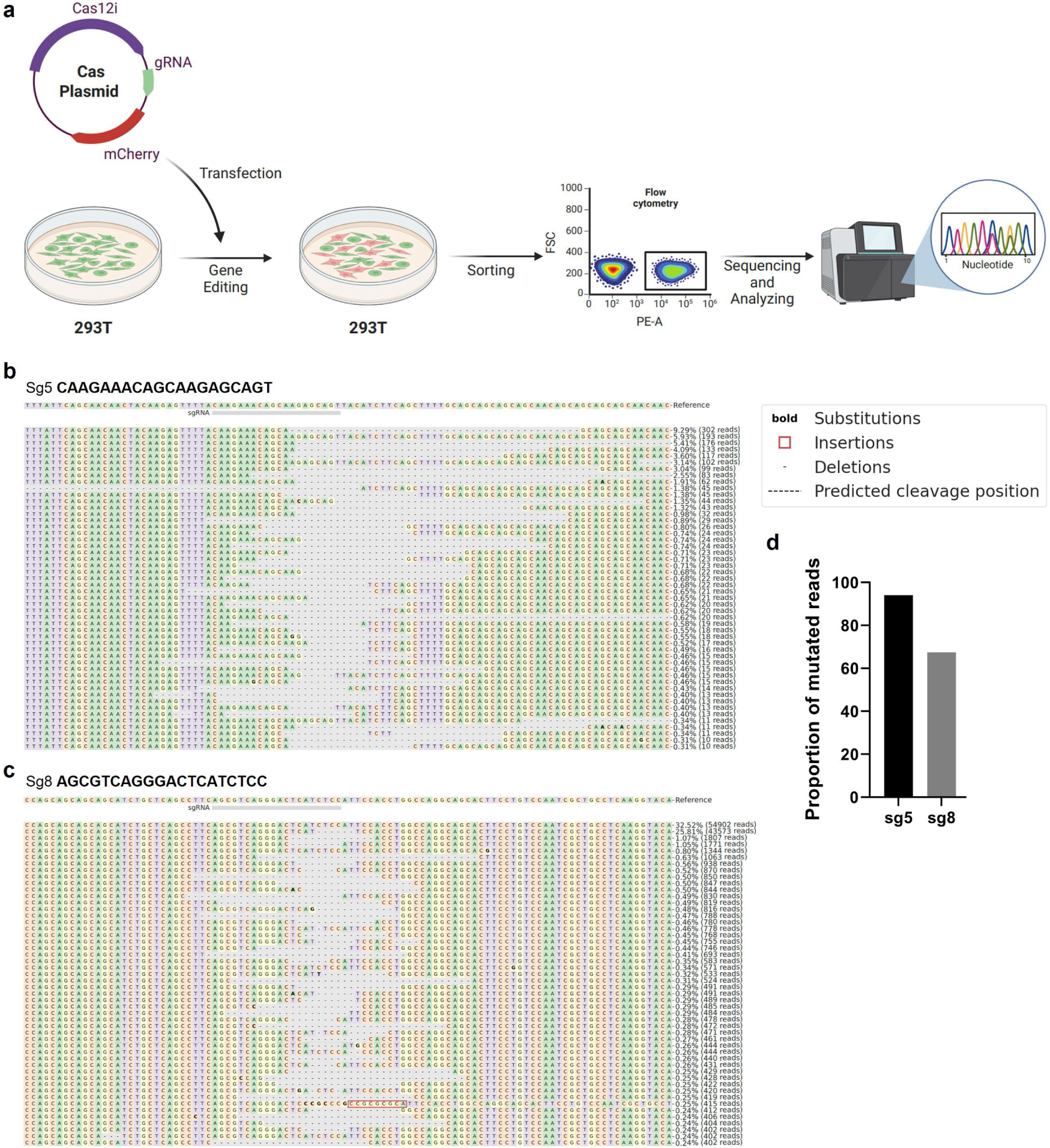
Screening of conserved sgRNAs targeting human and marmoset *FOXP2*. **a**, Schematic illustration of the sgRNA screening workflow. Candidate sgRNAs were selected based on high sequence identity between the human and marmoset genomic regions. A single-plasmid system expressing CRISPR-Cas components and an mCherry reporter was transfected into HEK293T cells via PEI. At 48 hours post-transfection, mCherry-positive cells were isolated by Fluorescence-Activated Cell Sorting (FACS). Genomic DNA was then extracted, and editing efficiency was evaluated using next-generation sequencing (NGS). **b-c**, Representative editing patterns for sgRNA5 (**b**) and sgRNA8 (**c**). The target sequences and their respective positions on the human reference genome are indicated. **d**, Histogram showing the percentage of mutated reads. Editing efficiency was quantified based on the frequency of insertions and deletions (indels).

**Extended Data Fig. 2.**
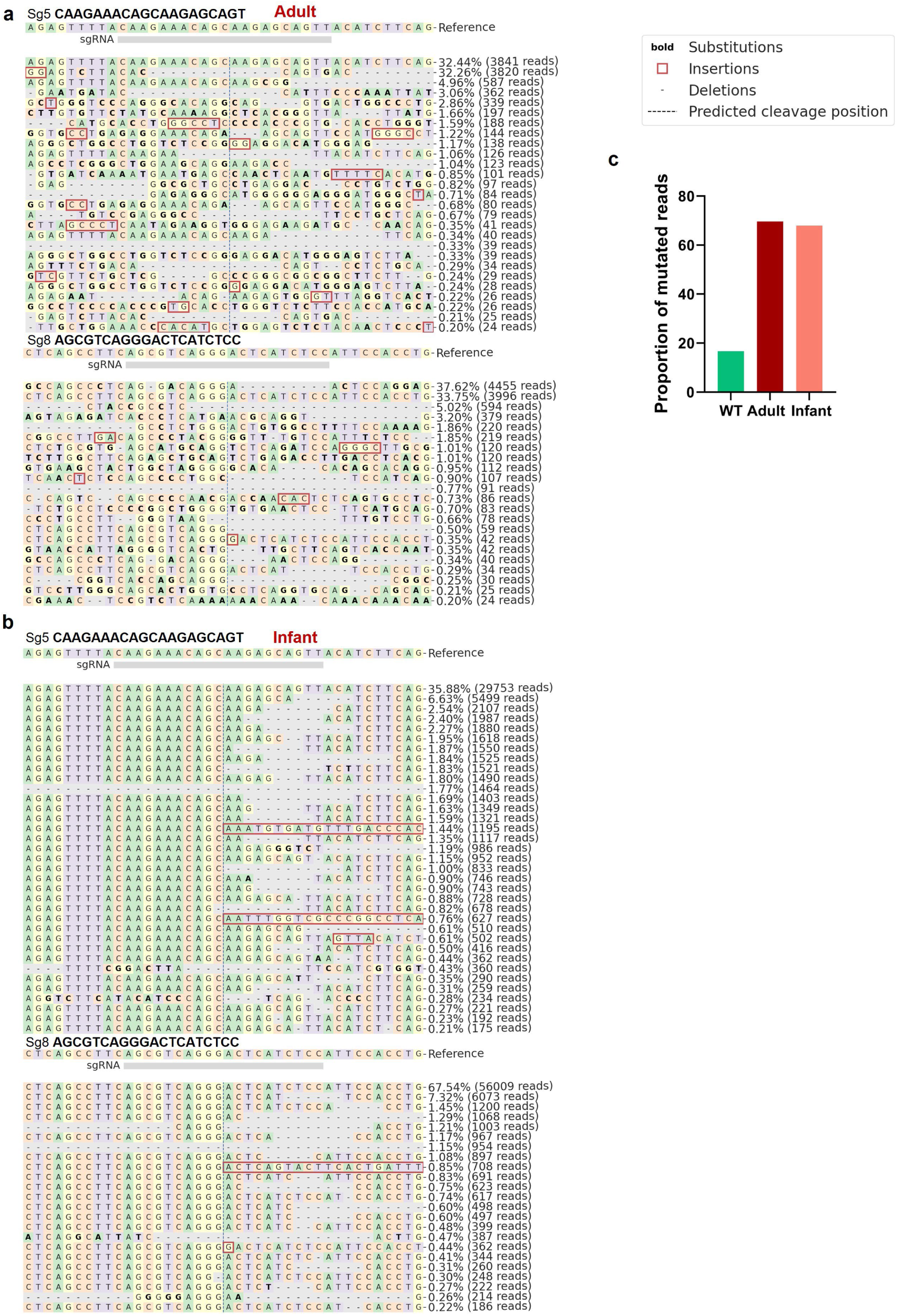
*In vivo* striatal *FOXP2* editing in adult and infant marmosets mediated by a single AAV vector carrying dual sgRNAs. **a-b**, Representative editing patterns at the sgRNA5 site (upper panels) and sgRNA8 site (lower panels) in an adult (**a**) and an infant (**b**) marmoset, respectively. Target sequences and their respective genomic coordinates on the marmoset reference genome (calJac3) are indicated. **c**, Histograms showing the percentage of mutated reads (indel frequency) at each target site. WT: wild type marmoset without AAV injection.

**Extended Data Fig. 3.**
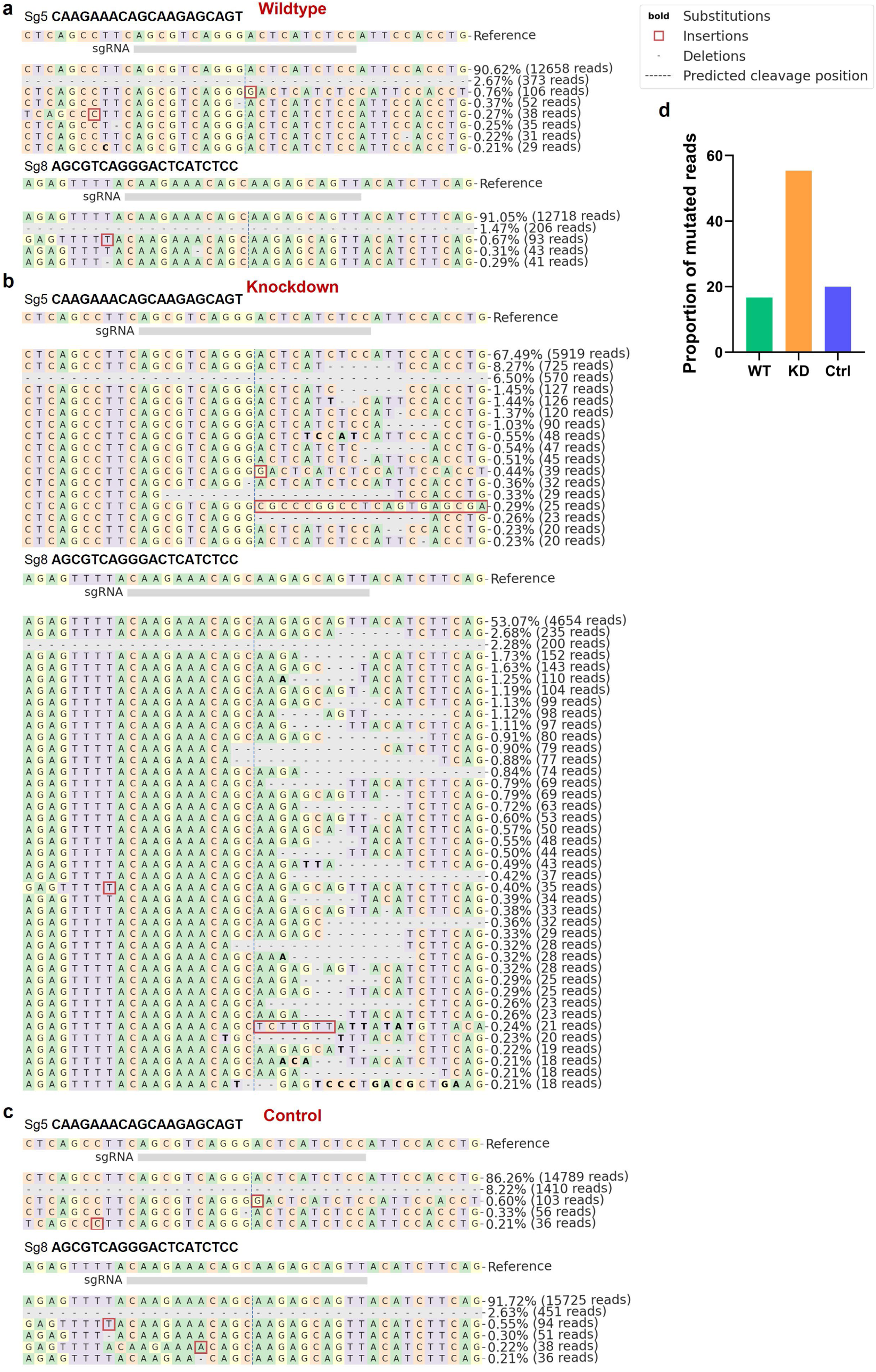
Validation of striatal *FOXP2* editing in recorded marmosets. **a-c**, Representative alignment plots showing editing patterns at the sgRNA5 site (upper panels) and sgRNA8 site (lower panels) in a wild type (WT) marmoset (**a**), a knockdown (KD) marmoset (**b**), and a control (Ctrl) marmoset (**c**). Target sequences and their respective genomic coordinates on the marmoset reference genome (calJac3) are indicated. **d**, Histograms representing the percentage of mutated reads (indel frequency) at each target site.

**Extended Data Fig. 4.**
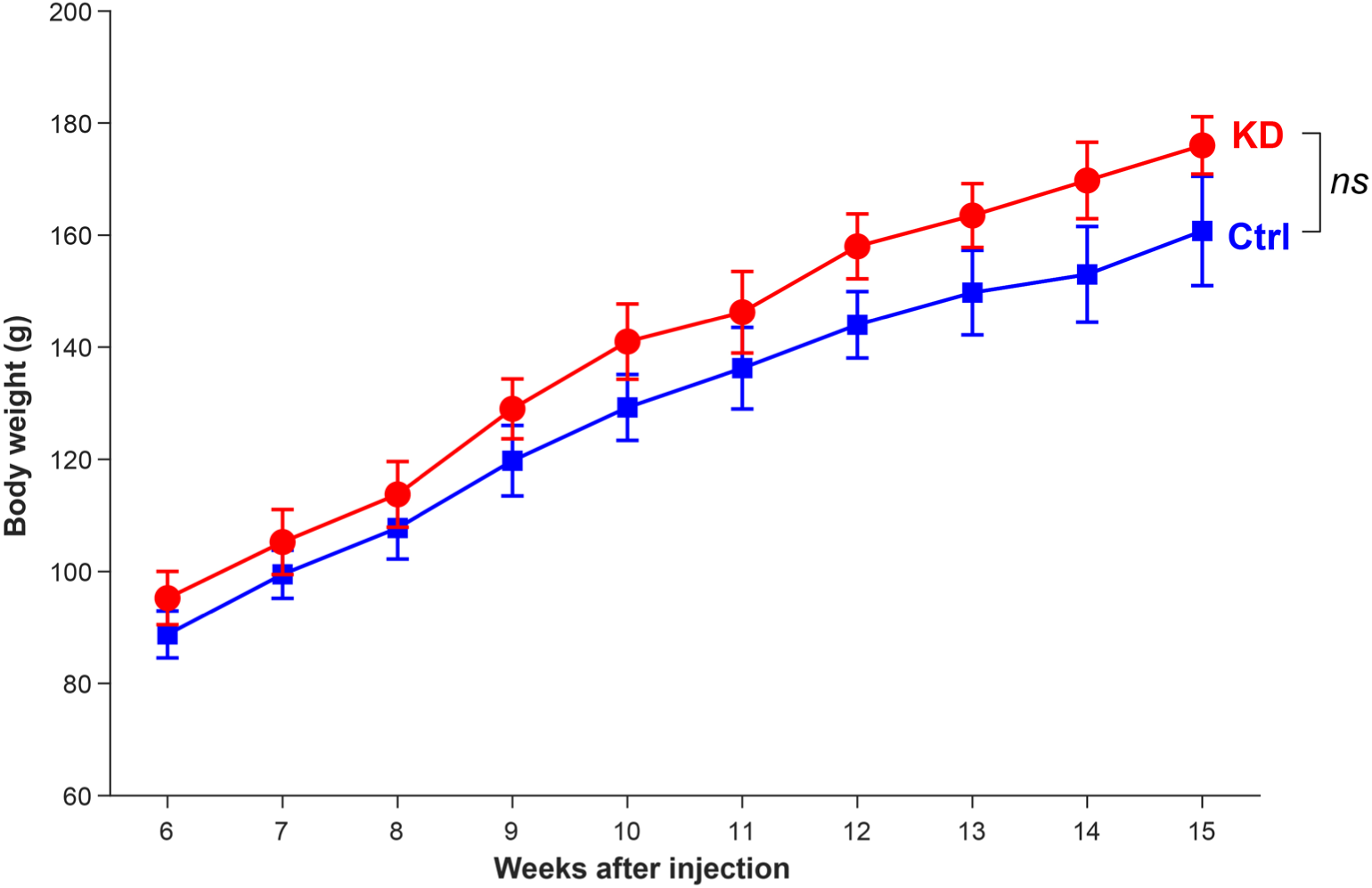
Body weights of marmosets in the FOXP2-edited and control groups. Body weights of marmosets in the control (Ctrl, *n* = 4) and knockdown (KD, *n* = 4) groups from week 6 to week 15 after virus injection. Data are mean ± s.e.m.; no significant differences in body weight were observed between the Ctrl and KD groups at any time point (paired t-test, all *P* > 0.05).

**Extended Data Fig. 5.**
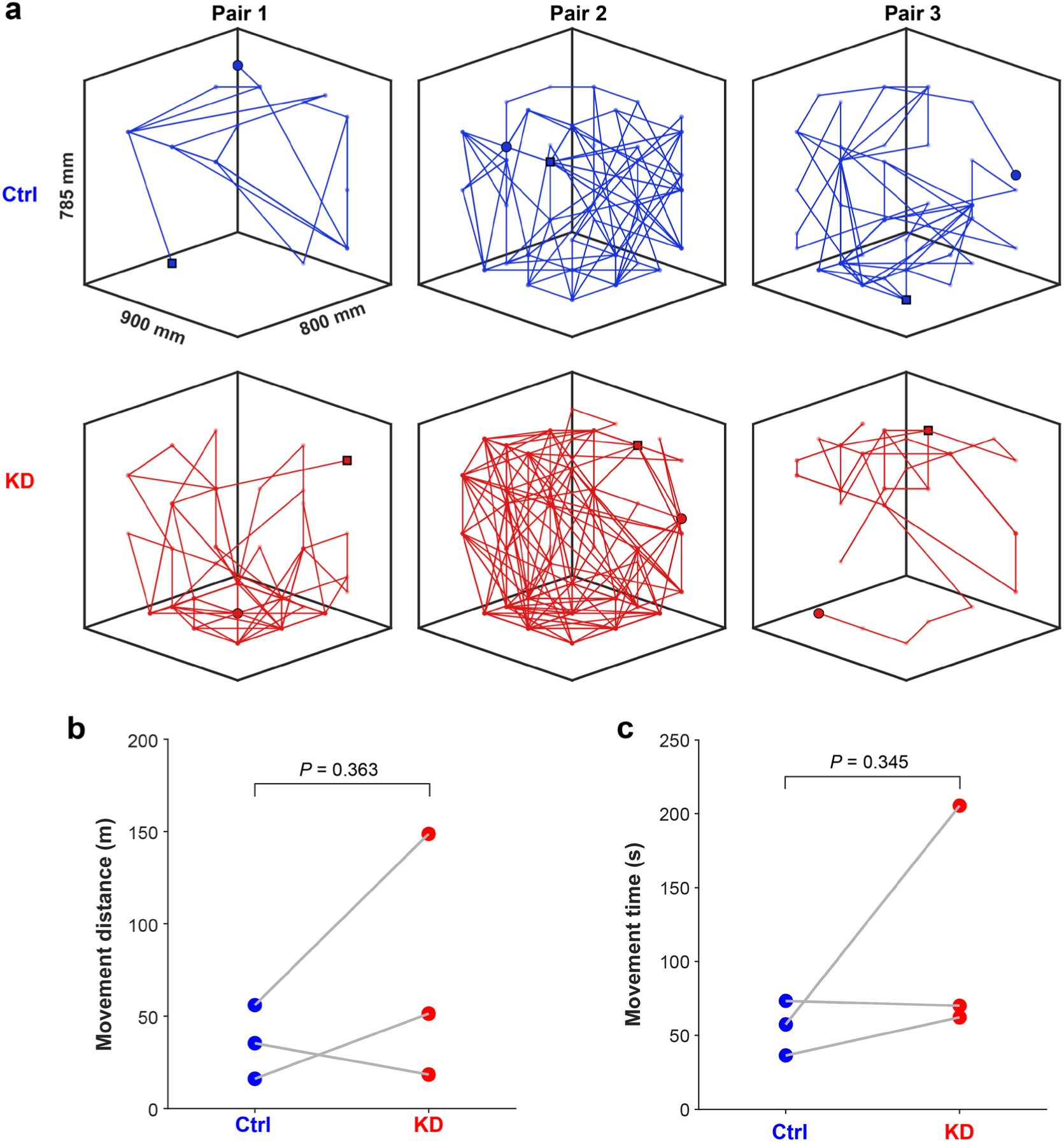
Global motion of marmosets in the FOXP2-edited and control groups. **a**, Motion trajectories of marmosets in the control (Ctrl, *n* = 3) and knockdown (KD, *n* = 3) groups during a 20-min video recording. Circles and squares indicate the start and end positions of each marmoset, respectively. **b**, Total distance traveled by each marmoset during the recording session. **c**, Total time spent in motion by each marmoset during the recording session. Statistical significance was assessed using paired t-test.

**Extended Data Fig. 6.**
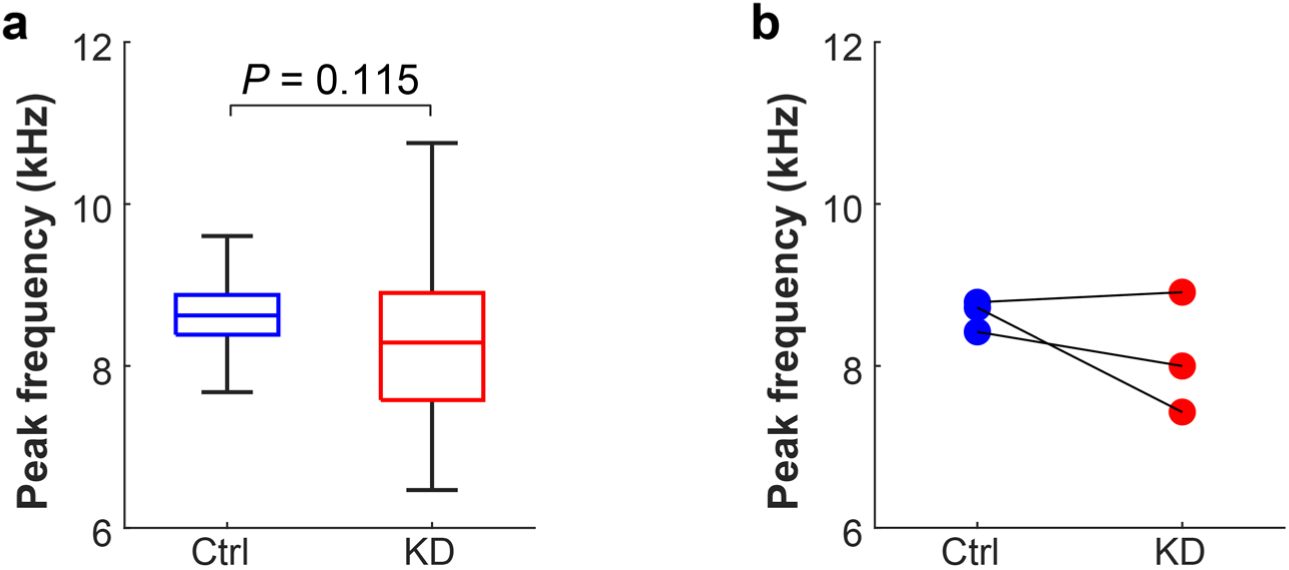
Comparison of peak frequency of phee syllables between marmosets in the FOXP2-edited and control groups. **a**, Peak frequency of phee syllables from marmosets in the control (Ctrl) and knockdown (KD) groups. Box plots indicate the 25th, 50th, and 75th percentiles, with whiskers extending to 1.5 × IQR. Statistical significance was assessed using linear mixed-effects models. **b,** Median peak frequency for each individual marmoset from the three pairs.

**Extended Data Table 1.**
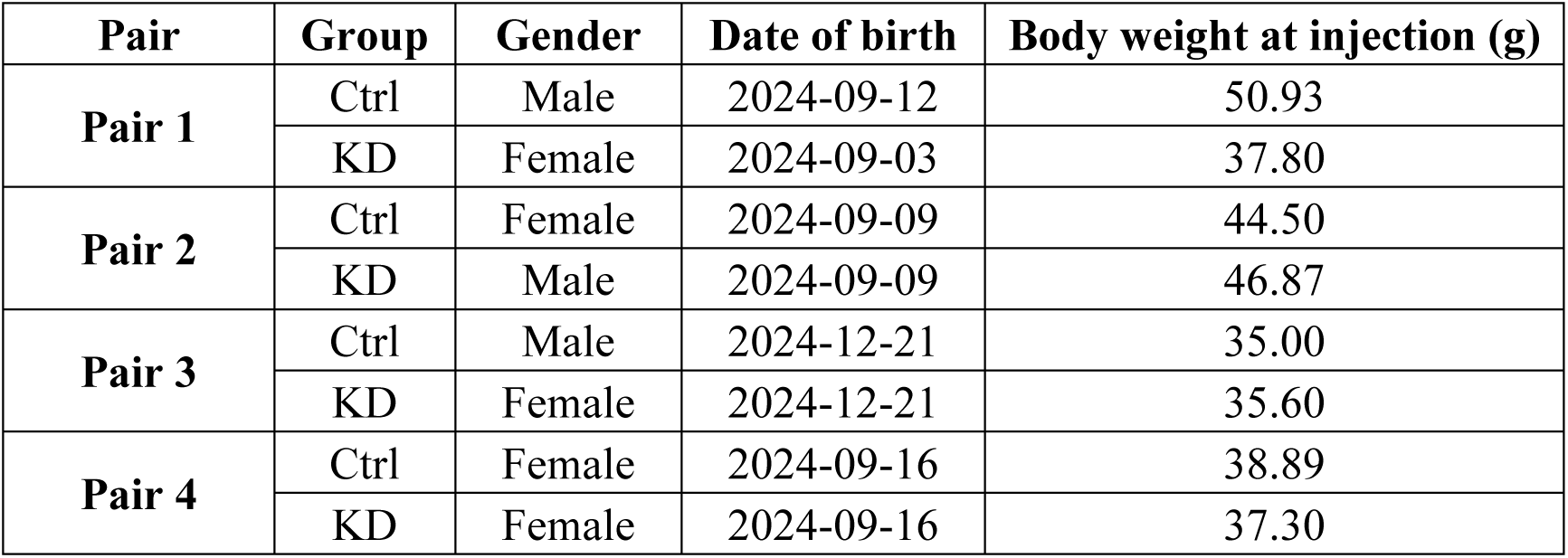
Characteristics of the four marmoset pairs.

